# Testing for the resistance of newly generated hybrid cacao germplasm in the gene pool of Cocoa Research Institute of Nigeria (CRIN) against *Phytophthora megakarya* pathogen causing black pod disease of cocoa

**DOI:** 10.1101/2020.12.07.414466

**Authors:** A. A. Tijani, A. H. Otuonye, M. O. Otusanya, A. O. Olaiya, O. O. Adenuga, C. G. Afolabi

## Abstract

Black pod disease caused by *P. Palmivora* and *P. megakarya* is an important disease affecting *cacao* in West Africa which produces 70% of the world output. Resistance to the pathogen is therefore, an important factor to be considered when breeding for high yielding and earliness in fruiting. Resistance to the pathogen using Leaf Disc Test (LDT) was conducted at the Plant Pathology Laboratory, Ibadan, on detached 15mm diameter discs of two-month-old leaves obtained from nineteen newly generated *cacao* hybrids in the gene pool of the Institute. Attached Whole Leaf Test (AWLT) was also conducted on few selected hybrids in the screen house at the same institute. Ten μl zoospores suspension of *P. megakarya* pathogen adjusted to inoculum strength of 3×10^5^ zoospores/ml using haemocytometer was used for the inoculation of the *cacao* LD. Using 0-5 disease rating scale, data was taken on the 5^th^ and 7^th^ day after inoculation for LDT and on the 6^th^ day after inoculation for AWLT. Data obtained were subjected to Analysis of Variance and significant means were separated using Student Newman Kuels Test at p<0.05. The LDT classified the hybrids into five groups namely: Resistant (hybrid 001, 003 and 005); Moderately Resistant (hybrid 006, 007, 008,013,015,018 and 019) Moderately Susceptible (hybrid 004, 014 and 017); Susceptible (hybrid 002, 010, 011 and 016) and Highly Susceptible (hybrid 009 and 012). Scores from LDT significantly correlated (r= 0.92**) with that of AWLT. It was observed from the study that *cacao* hybrid 001, 003 and 005 showed traits of resistant and therefore, could be incorporated into the national breeding programs for the development of high-yielding and resistant *cacao* cultivars. The moderately resistant hybrids could be conserved for future germplasm enhancement program.

## Introduction

Cacao (*Theobroma cacao* L.), an understory tree belongs to the order Malvales and family Malvaceae (1). It is an important cash crop cultivated for its nutritional value and industrial uses locally and internationally (2). Fourteen states in Nigeria namely Ondo, Cross River, Osun, Ogun, Ekiti, Edo, Abia, Adamawa, Taraba, Akwa Ibom, Delta, Kogi and Kwara were reported to produce 95% of country’s export (3). Reports showed that eight hundred thousand (800,000) hectares of Nigeria arable land were under *cacao* cultivation between 2003 – 2005 (4, 5). Cocoa production in Nigeria, due to several factors has failed to meet the demand from local and international consumption and processing. Prominent among these factors is the difficulty in the attainment of optimal yield from this tree crop despite the increased acreages largely due to cacao diseases.

Several pathogens attack *cacao*, but one of the most important diseases of *cacao* in Nigeria is the black pod disease caused by *Phytophthora* spp. Two of the several species of the black pod pathogen namely *P. palmivora* and *P. megakarya* have been identified as the causal agent of the disease in Nigeria. *P. palmivora* (less than 20% distribution) and *P. megakarya* (over 80% distribution) occur on *cacao* in Nigeria (3, 6–12). Of these two species reported on *cacao* in Nigeria, *P. megakarya* was noted to be more virulent (13). *P. megakarya* has also been reported to displaced *P. palmivora*, but the strategy involved is not clear (13).

In Nigeria, 30% yield and crop losses due to *P. palmivora* was reported (14) while 80% losses in *cacao* and its produce due to *P. megakarya* was reported in unkempt farms (15). In wetter part of the country, 90-100% yield losses were also recorded (16). Estimated annual yield losses of 700,000 metric tons of cocoa was reported globally due to black pod disease (17).Cultivation of susceptible varieties and the prevailing environmental conditions are factors that influence the disease (18).

Although chemical control is possible, it is expensive (Vis-a-vis 400kg/ha production) and require skills for effective application, couple with the Problem of chemical residue and environmental hazardousness (19–21). Screening for resistance to select cultivars that are resistant to the disease for field establishment has been advocated as the best disease management measure (22–25).

This management option is environmentally friendly and has the derivative advantage of been relatively cheap.

Therefore, this study was conducted to determine the resistance status of some newly generated cacao hybrids to black pod pathogen using leaf disc and attached whole leaf tests.

## Materials and methods

Nineteen newly produced cacao hybrids (Table 1) from a new cacao breeding research plot (7° 13’N, 3° 51’E) at Cocoa Research Institute of Nigeria (CRIN), Ibadan were used for the study in 2016.

**Table 1:** Cacao genotypes from hybrid gene pool selected for black pod disease resistance screening at CRIN

| Genotype | Hybrid | Pedigree |
| --- | --- | --- |
| (T82/27xT12/11) x (T101/15xN38) | 1 | Forastero |
| (T82/27xT12/11) x (T65/7xT22/28) | 2 | Forastero x (Amazon x Trinitario) |
| (T82/27xT12/11) x (T65/7xT57/22) | 3 | Forastero x (Amazon x Trinitario) |
| (T82/27xT12/11) x (T53/5xN38) | 4 | Forastero |
| (P<sub>7</sub>xT60/887) x (T65/7xT57/22) | 5 | Forastero x (Amazon x Trinitario) |
| (P<sub>7</sub>xT60/887) x (T53/5xN38) | 6 | Forastero |
| (T86/2xT9/15) x (T65/7xT57/22) | 7 | Amazon x Trinitario |
| (T86/2xT9/15) x (T53/5xN38) | 8 | (Amazon x Trinitario) x Forastero |
| (T86/2xT22/28) x (T65/7xT22/28) | 9 | Amazon x Trinitario |
| (T65/7xT9/15) x (T65/7xT22/28) | 10 | Amazon x Trinitario |
| (T65/7xT9/15) x (T65/7xT57/22) | 11 | Amazon x Trinitario |
| (T65/7xT9/15) x (T53/5xN38) | 12 | (Amazon x Trinitario) x Forastero |
| (P<sub>7</sub>xPa150) x (T101/15xN38) | 13 | Forastero |
| (P<sub>7</sub>xPa150) x (T65/7xT57/22) | 14 | Forastero x (Amazon x Trinitario) |
| (P<sub>7</sub>xPa150) x (T53/5xN38) | 15 | Forastero |
| 3-(P<sub>7</sub>xT60/887) x 5-(P<sub>7</sub>xT60/887) | 16 | Forastero |
| 3-(T65/7xT101/15) x 6-(T65/7xT101/15) | 17 | Forastero |
| 4-(T65/7xT101/15) x 6-(T65/7xT101/15) | 18 | Forastero |
| 6-(T65/7xT101/15) x 10-(T65/7xT101/15) | 19 | Forastero |
| C77 | 20 | Amazon |
| N38 | 21 | Amazon |
| Pa150 | 22 | Amazon |
| ICS1 | 23 | Amelonado |

Detached Leaf Discs Test (LDT) and Attached whole Leaf Test (AWLT) were employed for black pod resistance screening in CRIN plant pathology laboratory and greenhouse respectively. Genotype C77 and Pa150 were used as resistant checks while Genotype N38 and ICS1 were used as susceptible checks

### Procedure for Leaf Discs Test (LDT)

The cacao leaves used were obtained from the selected hybrid plants (Table 1) at CRIN headquarters, Ibadan. Healthy cacao leave of a similar age (approximately two-month-old) free of insect damage were harvested from the selected cacao hybrids early in the morning using secateurs within a space of two hours (07.00 - 09.00 am) to avoid the possible effect of harvesting time on the resistance level. A minimum of 5 leaves was randomly removed from the plants and kept in sterile polythene bags, moistened to maintain high relative humidity and then labeled properly and taken to the laboratory for study.

### Leaf Disc (LD) preparation

Adaxial and abaxial surfaces of the harvested leaves were cleaned with sterile towel to remove dirt and these were cut into 15 mm diameter discs with a semi-automated perforating device. The relative humidity in the incubation chamber was maintained at 80% by using a humidifier (Vicks VS100N Honeywell humidistat control) and cool air from a digital Samsung air conditioning unit to prevent the cut leaf discs from drying up.

The leaf discs were then distributed on a moistened absorbent paper towel in inoculation trays measuring 30 cm × 50 cm. The LD was arranged in the trays with the underside of the leaf facing upward. Ten leaf discs of each cacao hybrids including those of the four controls (C77, Pa150, ICS1 and N38) were distributed per tray, for the six trays that were used. There were 60 leaf discs in all for each progeny per inoculation series. This arrangement ensured that all the selected cacao hybrids were represented in each tray.

The cacao hybrids LD in the trays were numbered and sprayed with very fine droplets of sterile water from an atomizer (cat.No51901-spray unit Gellman Sciences, Ann, Arbor, Michigan, USA). This was to maintain the freshness of the discs up to the time of inoculation. These light-proof trays containing replicates of the cacao hybrids LD were covered and labeled to prevent mixed up. Each tray represents a block; hence, the arrangement forms randomize complete block design (RCBD). The trays were then placed in the incubation chamber.

### Identification of *Phytophthora* species

Cacao pods naturally infected by the *Phytophthora* species were harvested from the field. Then pod pieces were taken from the area of advancing lesion on the pod and were disinfected in 70 % ethanol and blotted dry with Whatman filtering paper No. 1. These pods pieces were then plated on a sterile potato dextrose agar (PDA) in 3 replicates. Subsequent transfers were made unto sterile PDA plates by hyphal tips to obtain pure cultures. Microscopic examination of slide cultures was made with x100 and 400 objectives of an Olympus compound microscope mounted with scope 9.0 digital imagery camera to identify relevant spore structures. Physical examination based on the characteristics “Seaweed” odour and culture pigmentation was done to identify the relevant species.

Pure cultures of the *Phytophthora* species were then stored for further uses.

### Inoculum Preparation

Pure cultures of the *Phytophthora* species obtained were inundated with 20ml of cold sterile water at 10°C. This was gently swirled and then the culture surface was scrapped with a sterile scalpel and filtered through 2-fold cheesecloth into a 50ml beaker to remove sporangia and agar debris.,It was further refrigerated at −5°C for 25 - 30 mins and then followed by incubation in the dark at 25°C±2°C for 30 - 45 minutes (26). The spore suspension was removed and kept at room temperature for one hour to decyst and then later decanted into an atomizer after it was adjusted to inoculum strength of 1 ×10^5^ zoospores/ml using a Malassez haemocytometer (France). 2mls of the spore suspension was used to spray the surface of each detached matured but unripe N38 genotype cacao pods in a plastic bowel. Lids were placed over the bowels and the pods were incubated at 25°C±2°C for 5 days to obtain sporulation and to ensure pathogenic virulence of the *Phytophthora* isolate used for the screening assay.

Zoospore was harvested by using the tip of soft brush to gently brush spores on artificially inoculated detached green *cacao* pod into a glass beaker into moderate quantity of 10°C sterile water and then zoospore suspension was prepared as described above. The release of motile zoospores (infective propagules) resulted from the rupturing of matured sporangia caused by alternating chilling and thawing of the suspension. The zoospore suspension was adjusted to inoculum strength of 3 ×10^5^ zoospores/ml using a Malassez haemocytometer (France).

### Leaf Disc Inoculation

The LD was inoculated the same day on which the leaf discs were prepared later in the afternoon. Inoculation was generally carried out by using sterile automated repeatable dispenser attached to a micro-syringe (Eppendorf). This was used to display 10μl droplets, discharge of the zoospore suspension in the middle of each leaf discs, at a right angle to the direction in which the discs of the genotypes were displayed in the trays. The light-proof trays were sealed hermetically and placed in a humidified chamber at air temperature of 26 ± 2°C and relative humidity 80-100% to incubate.The zoospore suspension was continuingly agitated prior and during the inoculation process to ensure even distribution of the zoospores.

The trays were inspected for lesion appearance, 72 hours after inoculation and data were collected for leaf discs reaction to black pod pathogen. Inoculation droplets that have not been absorbed yet were dried with filter paper and the trays were closed again. Subsequent observations were made on the 5th and 7^th^day of inoculation to described levels of infection of the LD using five-points disease assessment (rating) scale; 0-No symptoms, 1-small localized brown or dark-brown penetrations points, 2-small penetration points with some connection between them and small expanding lesion, 3-Coalescence of brown spots forming intermediate size lesions, 4-Expanding lesions, 5-Uniformly dark-brown large expanding lesions (38).

### Attached Whole Leaf Test

Attached leaves that correspond to about two months old leaves were inoculated with zoospores suspension (prepared and standardised as described for LDT) on selected *cacao* seedlings in the greenhouse. Eight (8) hybrids were randomly selected such that at least one hybrid represents each resistance class and replicated three (3) times. Two attached leaves were inoculated on each seedling which amounts to six (6) leaves per genotype for the eight (8) hybrids. Ten (10) points were inoculated at the underside of each selected leaf along the midrib which amounts to sixty (60) inoculated points per hybrid. Inoculation was carried out by using a sterile automated repeatable dispenser attached to a micro-syringe (Eppendorf) to deliver 10μl droplets of the zoospore suspension on each inoculation point, at a right angle along the leaf midrib. The greenhouse was maintained at 26 ± 2°C air temperature provided by digital Samsung air conditioning unit and approximately 80% relative humidity was provided by transparent polythene bags for the 6-day incubation period.

The inoculated leaves were inspected for lesion appearance, 6th day after inoculation and data were recorded on their reaction to black pod pathogen using a five-point disease assessment scale as described above. Both the Leaf Discs Test (LDT) and Attached Whole Leaf Test (AWLT) were repeated twice.

### Statistical analysis

Data obtained on the frequency and spread of lesions in leaf discs and attached leaf tests were subjected to analysis of variance (ANOVA) using SAS (Statistical Analysis System) software. Significant means were separated using the Student Newman Kuels Test (SNK) at p<0.05. Means obtained from leaf disc scores (for selected lines) were correlated with that of attached whole leaf to confirm the reliability of leaf disc test at p<0.05.

## Results

### Identification of *Phytophthora* species

The visual examination of the cultured plates showed that the diameter of black pod pathogen colony, 3 days after plating was 19mm.The pigmentation of the *P. megakarya* colony in culture plates even as it ages was milky-white to dirty-white. The aerial mycelium is of the fairly uniform deep cotton-wool like colony with faint lobed to floral pattern and diffused advancing margin. The *P. megakarya* culture was also noted to have a characteristic seaweed odour typical of *Phytophthora* species. The cardinal temperature range for its existence is 11°C minimum, optimum 24-28°C, and maximum 30°C.

The microscopic features of the sporangia morphology of the *Phytophthora* isolate was a round base prominently papillae with narrow to medium length non-occluded pedicel type, typical of *Phytophthora megakarya*.

### Evaluation of LD of the cacao Hybrids for resistance to *P. megakarya* isolate(s)

The result of the study showed that the newly produced cacao hybrids reaction to *P. megakarya* isolate(s) varied significantly with respect to the various lesion sizes induce on LD’s, (Table 2). Mean disease severity quantified by the lesions on LDs of the various cacao hybrid showed that (P_7_xT60/887) x (T65/7xT57/22) has a lowest mean disease score of 1.22 followed by (T82/27xT12/11) x (T65/7xT57/22) with a disease mean score of 1.31 and then the resistant check genotype Pa150 with a mean score of 1.33. The N38 control cacao genotype however had the highest disease mean score of 2.27 followed by cacao hybrid (T65/7xT9/15) x (T53/5xN38) with disease mean score of 2.17 and then the ICS1 and cacao hybrid (T86/2xT22/28) x (T65/7xT22/28) with disease mean scores of 2.15 and 2.14 respectively. Analysis of variance (ANOVA) done for LD’s lesion scores in their reaction to the *P. megakarya* isolates, (Table 2), shows that the resistance of newly generated cacao hybrids varied significantly at p<0.05. Significant means were separated using Student Newman Kuels (SNK) Test and classified as shown in Table 2.

**Table 2:** Mean lesion score of black pod disease induced by *Phytophthora* spp on the cacao hybrid leaf disc and their degree of resistance

| Genotype | Hybrid | Disease mean score | Resistance status |
| --- | --- | --- | --- |
| (T82/27xT12/11) x (T101/15xN38) | 1 | 1.3611<sup>ab</sup> | Resistant |
| (T82/27xT12/11) x (T65/7xT22/28) | 2 | 1.7722<sup>bc</sup> | Moderately susceptible |
| (T82/27xT12/11) x (T65/7xT57/22) | 3 | 1.3056<sup>a</sup> | Resistant |
| (T82/27xT12/11) x (T53/5xN38) | 4 | 1.8889<sup>cde</sup> | Susceptible |
| (P<sub>7</sub>xT60/887) x (T65/7xT57/22) | 5 | 1.2722<sup>a</sup> | Resistant |
| (P<sub>7</sub>xT60/887) x (T53/5xN38) | 6 | 1.6611<sup>abc</sup> | Moderately resistant |
| (T86/2xT9/15) x (T65/7xT57/22) | 7 | 1.6333<sup>abc</sup> | Moderately resistant |
| (T86/2xT9/15) x (T53/5xN38) | 8 | 1.6722<sup>abc</sup> | Moderately resistant |
| (T86/2xT22/28) x (T65/7xT22/28) | 9 | 2.1444<sup>de</sup> | Highly susceptible |
| (T65/7xT9/15) x (T65/7xT22/28) | 10 | 1.7944<sup>bcd</sup> | Moderately susceptible |
| (T65/7xT9/15) x (T65/7xT57/22) | 11 | 1.8389<sup>bcd</sup> | Moderately susceptible |
| (T65/7xT9/15) x (T53/5xN38) | 12 | 2.1667<sup>de</sup> | Highly susceptible |
| (P<sub>7</sub>xPa150) x (T101/15xN38) | 13 | 1.6389<sup>abc</sup> | Moderately resistant |
| (P<sub>7</sub>xPa150) x (T65/7xT57/22) | 14 | 1.9722<sup>cde</sup> | Susceptible |
| (P<sub>7</sub>xPa150) x (T53/5xN38) | 15 | 1.6778<sup>abc</sup> | Moderately resistant |
| 3-(P<sub>7</sub>xT60/887) x 5-(P<sub>7</sub>xT60/887) | 16 | 1.8222<sup>bcd</sup> | Moderately susceptible |
| 3-(T65/7xT101/15) x 6-(T65/7xT101/15) | 17 | 1.9500<sup>cde</sup> | Susceptible |
| 4-(T65/7xT101/15) x 6-(T65/7xT101/15) | 18 | 1.5778<sup>abc</sup> | Moderately resistant |
| 6-(T65/7xT101/15) x 10-(T65/7xT101/15) | 19 | 1.5389<sup>abc</sup> | Moderately resistant |
| C77 | 20 | 1.8389<sup>bcd</sup> | Moderately susceptible |
| N38 | 21 | 2.2667<sup>e</sup> | Highly susceptible |
| Pa150 | 22 | 1.3389<sup>a</sup> | Resistant |
| ICS1 | 23 | 2.1500<sup>de</sup> | Highly susceptible |

Means in the same column not followed by the same letter are significantly different according to Student Newman Kuels (SNK) Test at 5% probability level

At p<0.05, there was no significant difference in the reactions of cacao hybrid (T82/27xT12/11) x (T65/7xT57/22), (P7xT60/887) x (T65/7xT57/22) and the resistant check cacao genotype Pa150 with the *P. megakarya* isolates and are classified as resistant, Table 2. The cacao hybrid (T82/27xT12/11) x (T101/15xN38), react significantly different but closer to others in resistant category, thus classified as resistant. However, the reaction of these cacao hybrids and genotype to the *P. megakarya* isolates significantly varied with the other cacao hybrids and genotypes. Cacao hybrids (P_7_xT60/887) x (T53/5xN38), (T86/2xT9/15) x (T65/7xT57/22), (T86/2xT9/15) x (T53/5xN38), (P_7_xPa150) x (T101/15xN38), (P_7_xPa150) x (T53/5xN38), 4-(T65/7xT101/15) x 6-(T65/7xT101/15) and 6-(T65/7xT101/15) x 10-(T65/7xT101/15) did not differ either in their reaction to the *P. megakarya* isolates but varied significantly in their reaction when contrasted with the other cacao hybrids and genotypes and hence classified to be moderately resistant. The other cacao hybrids and genotypes reaction to the *P. megakarya* isolates varied from moderate to been highly susceptible.

In the attached leave assay, Pa150 has the lowest mean lesion score of 0.95 followed by cacao hybrid (P_7_xT60/887) x (T65/7xT57/22) and (T82/27xT12/11) x (T101/15xN38) with mean lesion score of 1.14 and 1.15 respectively. N38 cacao genotype had the highest mean lesion score of 1.94 followed by cacao hybrid (T86/2xT22/28) x (T65/7xT22/28) with 1.84 and then ICS1 cacao genotype with lesion mean score of 1.71 (Table 3). ANOVA ran for disease mean score on attached leaves showed significant variations in the various reactions of the cacao hybrids and the check (control) genotypes. At p<0.05, Pa150 significantly varied in its reaction with the other cacao hybrids and genotypes and was classified as highly resistant. Cacao hybrid (P_7_xT60/887) x (T65/7xT57/22) and (T82/27xT12/11) x (T101/15xN38) did not vary in their reaction to the *P. megakarya* isolates but significantly varied in their reaction to *P. megakarya* isolates when compared with the other cacao hybrids and genotypes and are classified as resistant, (Table 3). Cacao hybrid (T86/2xT9/15) x (T65/7xT57/22) also differed significantly in its reaction with the other cacao hybrids and genotypes and was classified as moderately resistant. The other cacao genotypes and hybrids’ reaction to *P. megakarya* showed various reactions that ranged from moderately susceptible to susceptible.

**Table 3:**
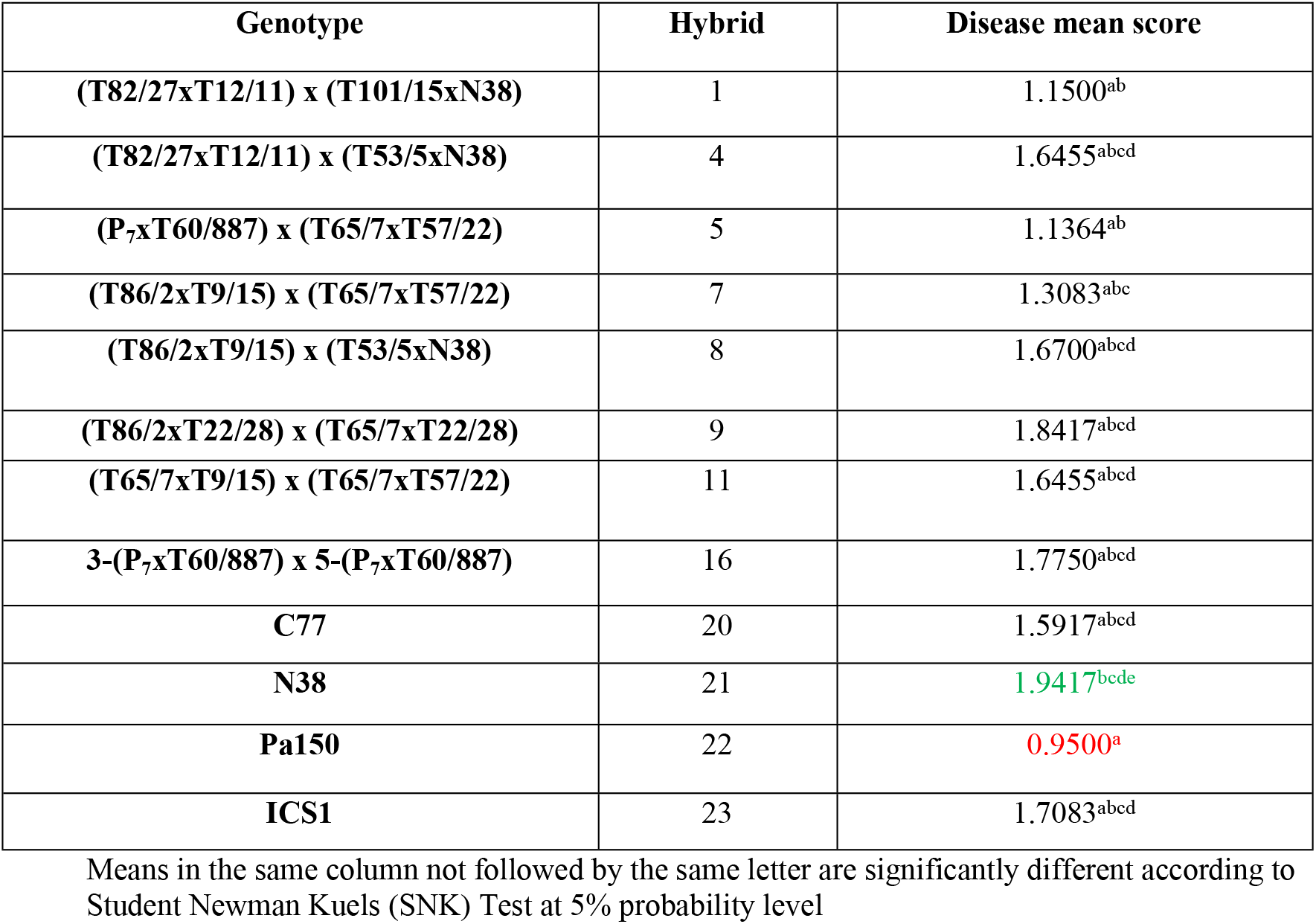
Mean lesion score of black pod disease induced by *Phytophthora* spp. on the cacao hybrid attached whole leaf inoculation (selected hybrid)

A good correlation was found between leaf discs and attached whole leaf score means at p<0.05 (r= 0.93**), Table 4.

**Table 4:**
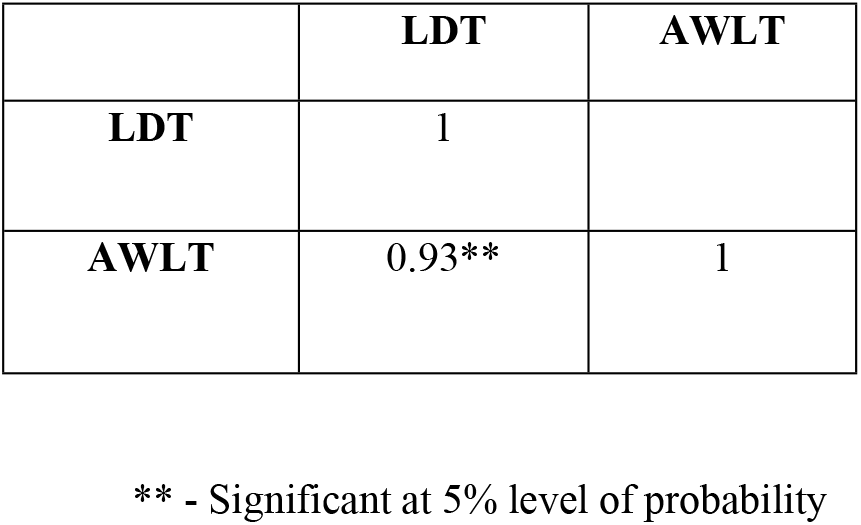
Correlation of Leaf Disc Test (LDT) with Attach Whole Leaf Test (AWLT)

|  | LDT | AWLT |
| --- | --- | --- |
| LDT | 1 |  |
| AWLT | 0.93** | 1 |

\*\* - Significant at 5% level of probability

## Discussion

The study shows the intrinsic variable genetic qualities existing among cacao hybrids assayed for resistance to black pod disease in the gene pool of CRIN. The mean lesion score of the disease severity in the detached LD assay of the cacao hybrids showed that none of the hybrids was totally immune to the black pod pathogen. This observation is in agreement with earlier reports of some authors (26, 27).

It was also observed in the detached LD assay in the study that the cacao hybrids (T82/27xT12/11) x (T65/7xT57/22) and (P7xT60/887) x (T65/7xT57/22) showed resistance to the *P. megakarya* pathogen which was not significantly different from the resistant control Parinary 150 (Pa150) that is of the Forastero group and an Upper Amazonian cacao genotype. This finding is in agreement with the reports of some authors (24, 28) who reported that the Forasteros groups, which include the Amazon cacao genotypes, are made up of higher percentage of moderately resistant to resistant genotypes. It was also noted that Trinitarios, been hybrid progeny between Criollo and Forastero are highly heterogeneous in their resistant character (24, 28). This may be responsible for the observed reactions of the cacao hybrids to *P. megakarya* black pod pathogen.

The study also showed that seven (7) cacao hybrids with Forastero, Amazon and Trinitario pedigree showed moderate resistance to the *P. megakarya* pathogen. This conformed to the findings of Iwaro *et al*. and Otuonye. It was also reported that cacao genotype T12/11 and T86/2, both Upper Amazon cacao genotypes, crossed to obtain parents for some of the hybrids that showed resistant and moderate resistant respectively to *P. megakarya* pathogen (24, 28). Reports from other workers (2, 30, 31) showed that T12/11 and T86/2, Sca6, T60/887, Pa 150, P7 and Pa 7/808 were used as crosses for some of the parents of the hybrids and resistant control as black pod disease resistant/escaping clones. In Cameroon, highest resistance to *Phytophthora* pod rot was also reported for hybrid P7 x Pa150 using LD assay which was correlated with highest resistance obtained in the field with this cacao genotype (32, 33).

It was observed from the detached LD assay in this study that nine (9) of the cacao hybrids with Forastero, Amazon and Trinitario pedigree showed moderate susceptibility to high susceptibility class with none of the cacao hybrids however, showing worst reaction when compared with the susceptible control ICS1 and N38 genotypes. The observation is in agreement with the findings of some scientists (28, 34–36) who reported that variation in character expression in cacao progenies given the Quantitative trait loci (QTL) of the parents of a progeny may involve either of the parents in character expression if they favorably share a common alleles brought by both or the character expression may involve each individual parent where it is not shared. They also noted that the variation in character expression between and within clones and progeny families may be due to segregation of the genome in response to resistance. Forastero is a very variable group and consist of the lower Amazon (West African ‘Amelonado’, ‘Maranhoa’, ‘Comun’ and ‘Para’ types from Brazil) and the Upper Amazonian cocoas (37). The lower Amazonian cacao genotypes for example, the West African ‘Amelonado’ are known for their susceptibility to the black pod disease. The out-breeding nature of cacao which is responsible for a high level of heterogeneity may bring about resistant variation observed in Trinitarios, been a hybrid between Forastero and Criollo (24). The variation in character expression by these cacao hybrids may be due to an involvement and expression of alleles of an individual of the parents in the QTL. This may account for the variation observed in the resistant character expressed by these cacao hybrids to the *P. megakarya* pathogen.

In the attached whole leaf assay, a high resistant trait was expressed by Pa150 in the study confirming its reliability as a resistant check. This observation was same for all the cacao hybrids and control genotypes selected for the attached whole leaf assay. It was observed for example that cacao hybrid (T82/27xT12/11) x (T53/5xN38) that showed susceptibility in the detached LD assay showed moderate susceptibility in attached whole leaf assay. This finding is in conformity with the findings of Iwaro *et al*. and Otuonye (24, 28) who reported that susceptibility is generally higher in detached leaves and pods than in the attached organs. This may be responsible for the observed reaction of the attached whole leaf of the cacao hybrids to the *P. megakarya* pathogen.

The insignificant isolate x host interaction has been reported from the findings by several scientists (2, 25, 38-40) established *Phytophthora* species non-specificity with cacao genotypes. This indicates that cacao hybrids resistance in this study to *P. megakarya* could be applicable to other *Phytophthora* species causing pod rot of cacao in the other areas of cocoa production in the world.

It was observed also from the study that C77 cacao genotype which was one of the resistant controls (28, 29) showed moderately susceptibility to the *P. megakarya* pathogen. This finding points to the fact that there may be changes in the population of the *P. megakarya* pathogen on cacao in Nigeria which indicates possible diversity in the genetic make-up of the pathogen with the appearance of more virulent strains that enable it to breakdown the resistance of C77 cacao genotype and cause enormous losses in revenue for the cocoa industry. This point to the need for continuous screening of cacao germplasm with the leaf disc assay, which have been proven to be reliable, cheap and fast in the selection and enhancement of germplasm for resistance to black pod disease.

A good correlation obtained between detached leaf disc test (LDT) and the attached whole leaf test (AWLT) in this study showed the reliability of the detached LD assay as an effective screening method that could be used to predict cacao genotypes reaction and resistance in the field to *P. megakarya* pathogen.

## Conclusion

- This study revealed that some of the cacao hybrids used in this study had innate resistant trait against *Phytophthora* pathogen, the causal pathogen of black pod disease of cacao, in the gene pool of CRIN. Of the nineteen cacao hybrids screened, three (3) were found to be resistant, seven (7) were found to be moderately resistant, four (4) were found to be moderately susceptible, three (3) were found to be susceptible and two (2) were found to be highly susceptible.
- C77 cacao genotype which was previously reported and consequently use as resistant check was found to be moderately resistant in this study, thus appearing to be losing its hitherto resistant status.
- The resistance in attached leaves of cacao hybrids is higher than that of the detached leaves and the reliability of Pa150 as resistant check remained intact.

## Recommendation

Cacao hybrid (T82/27xT12/11) x (T101/15xN38), (T82/27xT12/11) x (T65/7xT57/22) and (P_7_xT60/887) x (T65/7xT57/22) which were classified to be resistant in this study, are suggested for selection in cacao breeding activities for resistant varieties. These can be recommended for incorporation into national breeding program for distribution and developing high yielding disease resistant cacao cultivar. While hybrid [(P_7_xT60/887) x (T53/5xN38), (T86/2xT9/15) x (T65/7xT57/22), (T86/2xT9/15) x (T53/5xN38), (P_7_xPa 150) x (T101/15xN38), (P_7_xPa 150) x (T53/5xN38), 4-(T65/7xT101/15) x 6-(T65/7xT101/15) and 6-(T65/7xT101/15) x 10-(T65/7xT101/15) which were found to be moderately resistant could be conserved for future germplasm enhancement program.

